# The transcriptional landscape of cortical interneurons underlies *in-vivo* brain function and schizophrenia risk

**DOI:** 10.1101/481036

**Authors:** Kevin M Anderson, Meghan A Collins, Rowena Chin, Tian Ge, Monica D Rosenberg, Avram J Holmes

**Affiliations:** Department of Psychology, Yale University, New Haven, Connecticut 06520; Psychiatric and Neurodevelopmental Genetics Unit, Center for Genomic Medicine, Massachusetts General Hospital, Boston, MA 02114, USA; Department of Psychiatry, Massachusetts General Hospital, Harvard Medical School, Boston, MA 02114, USA; Department of Psychiatry, Yale University, New Haven, Connecticut 06520, USA

**Keywords:** fMRI, interneuron, genetics, somatostatin, parvalbumin, schizophrenia

## Abstract

Inhibitory interneurons orchestrate information flow across cortex and are implicated in psychiatric illness. Although classes of interneurons have unique functional properties and spatial distributions throughout the brain, the relative influence of interneuron subtypes on brain function, cortical specialization, and illness risk remains elusive. Here, we demonstrate stereotyped organizational properties of somatostatin and parvalbumin related transcripts within human and non-human primates. Interneuron spatial distributions recapitulate cortico-striato-thalamic functional networks and track regional differences in functional MRI signal amplitude. In the general population (n=9,627), parvalbumin-linked genes account for an enriched proportion of genome-wide heritable variance in *in-vivo* functional MRI signal amplitude. This relationship is spatially dependent, following the topographic organization of parvalbumin expression in independent post-mortem brain tissue. Finally, genetic risk for schizophrenia is enriched among interneuron-linked genes and predictive of cortical signal amplitude in parvalbumin-biased regions. These data indicate that the molecular genetic basis of resting-state brain function across cortex is shaped by the spatial distribution of interneuron-related transcripts and underlies individual differences in risk for schizophrenia.

**Key Findings:**

1. Spatial distributions of somatostatin (*SST*) and parvalbumin (*PVALB*) are negatively correlated in mature human and non-human primate cortex, paralleling patterns observed *in utero*.
2. *SST* and *PVALB* are differentially expressed within distinct limbic and somato/motor cortico-striato-thalamic networks, respectively.
3. *In-vivo* resting-state signal amplitude is heritable in the general population and tracks relative *SST/PVALB* expression across cortex.
4. Single-nucleotide polymorphisms tied to *PVALB*-related genes account for an enriched proportion of the heritable variance in resting-state signal amplitude.
5. *PVALB*-mediated heritability of resting-state signal amplitude in the general population is spatially heterogeneous, mirroring the cortical expression of *PVALB* in independent post-mortem brain tissue.
6. Polygenic risk for schizophrenia is enriched among interneuron-linked genes and predicts resting-state signal amplitude in a manner that also follows the cortical expression of *PVALB*.

## Introduction

Ramón y Cajal theorized that the functional diversity of the human brain arises, in part, from the vast assortment of neurons that pattern cortex^1^. Inhibitory interneurons are the most varied neuronal class^2^, exhibiting divergent morphological and physiological properties and coordinating information flow across the brain’s collective set of functional connections (functional connectome)^3,4^. Foundational cross-species animal and human work provides converging evidence for the role of interneurons in healthy brain functions as well as their dysregulation in psychiatric illnesses, including schizophrenia^5,6^ and major depressive disorder^7^. The development of densely sampled gene transcriptional atlases now enables the study of cellular and molecular correlates of functional brain network architecture^8-11^. Despite these methodological advances and a clear role for interneurons in the modulation of excitatory neuron activity, relatively little is known about how the spatial distribution of interneuron subtypes shape human brain activity and associated risk for psychiatric illness.

The topographic distribution of interneuron subtypes is theorized to contribute to regional and functional network specialization, partly by altering the relative excitatory/inhibitory balance within a given patch of cortex^9,12,13^. Interneurons comprise approximately 20-30% of cortical neurons^14^ and form stereotyped microcircuits with excitatory projection neurons^15^. While the precise number of interneuron subtypes is under debate, the vast majority express one of a limited set of genetic markers: somatostatin (*SST*), parvalbumin (*PVALB*), and vasoactive-intestinal peptide (*VIP;* a subset of *HTR3A* interneurons)^2^. Each molecular subtype possesses unique synaptic and functional characteristics, leading to the hypothesis that the ratio of specific interneuron classes may drive local differences in neural activity. For example, *SST* expressing interneurons preferentially target dendrites of cortical projection neurons to regulate input whereas *PVALB* expressing interneurons primarily synapse on perisomatic regions to regulate output^2,16^. Consequently, the increased presence of SST, relative to other classes of interneurons, may facilitate filtering of noisy or task-irrelevant cortical signals as well as increase recurrent excitation required for higher-order cognition^20^. Conversely, relative increases in PVALB may produce stronger feedback inhibition on excitatory neurons^13^, leading to shorter activation timescales^17^ suited for processing constantly changing sensorimotor stimuli. These collective results suggest that the spatial distribution of interneuron subtypes could underlie regional differences in temporal signaling across cortex, as indexed by blood oxygenation level-dependent (BOLD) functional magnetic resonance imaging (fMRI).

Establishing the organizational principles by which cellular diversity influences brain function is a long-standing challenge in neuroscience, and could provide a route to understand individual variability in the diverse processing capabilities of the human brain across health and disease. Consistent with this aim, recent translational work suggests a core role of PVALB interneurons in the biological basis of fMRI measures of *in-vivo* brain function^18^. PVALB interneurons are known to orchestrate gamma-band oscillations (30-80 Hz^19,20^), a frequency range that is tightly coupled to spontaneous BOLD fluctuations^21-25^. Optogenetic stimulation of PVALB interneurons in rodents drives rhythms in the gamma range, impacting information processing through the synchronization of excitatory neurons^20^. In psychiatric illness, several lines of evidence suggest that decreased PVALB-mediated inhibition may serve as a core locus of disruption in schizophrenia, giving rise to the altered gamma-band signal and working memory deficits observed in the disorder^26^. However, a direct link between PVALB-related genetic variation and *in-vivo* brain activity has yet to be established. Linking cortical interneurons to individual differences in human brain function would yield deep insight into the biological basis of the hemodynamic BOLD signal, providing an engine for the discovery of functional connectome-linked genes and associated risk for illness onset.

Here, we bridge genetic, transcriptional, and neuroimaging data to advance three related lines of inquiry linking interneurons to human brain function. First, we describe the principal organizational features of *SST* and *PVALB* expression in both human and non-human primates, demonstrating a robust pattern of anti-correlation across cortex. Supporting the hypothesis that interneuron ratios contribute to functional specialization, *SST* and *PVALB* were differentially expressed within distinct limbic and somato/motor cortico-striato-thalamic functional loops, respectively. Second, we establish that the relative density of *SST* and *PVALB* tracks regional differences in brain activity across cortex. In a population-based sample of 9,627 individuals^27^, genetic variation among *PVALB*-correlated genes accounted for an enriched proportion of heritable variance in resting-state signal amplitude in a manner that mirrors the spatial expression of *PVALB* in an independent analysis of post-mortem brain tissue. Critically, these discoveries suggest that the molecular genetic basis of cortical function is not spatially uniform and that genes linked to PVALB interneurons underlie heritable aspects of the BOLD response. Third, we find evidence supporting the link between PVALB interneurons and psychotic illness, demonstrating that genetic risk for schizophrenia is enriched among interneuron-linked genes while also predicting reduced resting-state signal amplitude in a spatially heterogenous manner that follows the cortical expression of *PVALB.* These data help to address a long-standing challenge of neuroscience to understand how cytoarchitecture shapes human brain function and related vulnerability for psychiatric illness.

## Results

### Stereotyped anti-correlation of SST and PVALB interneuron markers across cortex

The unique properties of interneuron subtypes emerge early in development and are determined, in part, by their spatial location of origin in the embryonic ganglionic eminence^28,29^. *VIP* interneurons are born within the caudal ganglionic eminence (CGE), whereas *SST* and *PVALB* interneurons originate in the medial ganglionic eminence (MGE) along negatively correlated spatial gradients^30^. Parvalbumin- and somatostatin-destined neurons differentially cluster within the dorsal and ventral MGE, respectively^31,32^. Evidence in humans^9,12^ and rodents^13^ indicates that *SST* and *PVALB* maintain a negative spatial correlation in adulthood, suggesting that embryonic gradients may constitute a “proto-map” of mature cortex. Although the functional consequences of a negative spatial *SST/PVALB* relationship are not well understood, the presence of replicable and evolutionarily conserved expression patterns may suggest the importance of such interneuron gradients.

To characterize interneuron topography across human and non-human primate cortex, we analyze gene expression data from the Allen Human Brain Atlas (AHBA)^33^ and NIH Blueprint Non-Human Primate (NHP) Atlas ^34^. Cortical tissue AHBA samples from the left (*n*=1,273) and right (*n*=428) hemispheres were analyzed. Microarrays do not give absolute estimates of gene transcription, but can measure within-probe differences across samples. *SST* and *PVALB* expression values were mean and variance normalized across cortical samples, and subtracted (i.e. *SST*-*PVALB*) to reveal relative expression differences (Figure 1a). Extending prior evidence of negative spatial expression relationships between *SST* and *PVALB*^9,12^, these two transcripts were inversely correlated across available AHBA cortical samples (Figure 1e; *r*(1,699)=-0.43, p<2.2e-16). *SST* and *PVALB* distributions were organized along an anterior to posterior gradient, with relative *SST* expression greatest in orbitofrontal and medial prefrontal cortex, anterior insula, anterior cingulate, and the temporal lobe (Figure 1a-c; Supplemental Figure 1). In contrast, relative *PVALB* expression was greatest within unimodal sensory, motor, and visual cortices, as well as the parietal lobe. Histologically defined anatomical categories were used to characterize regional differences of interneuron density (Figure 1c). Median relative expression of *SST* and *PVALB* was negatively correlated across cortical subregions (*r*(39)*=-*0.87, p=1.0e-13).

**Fig 1.**
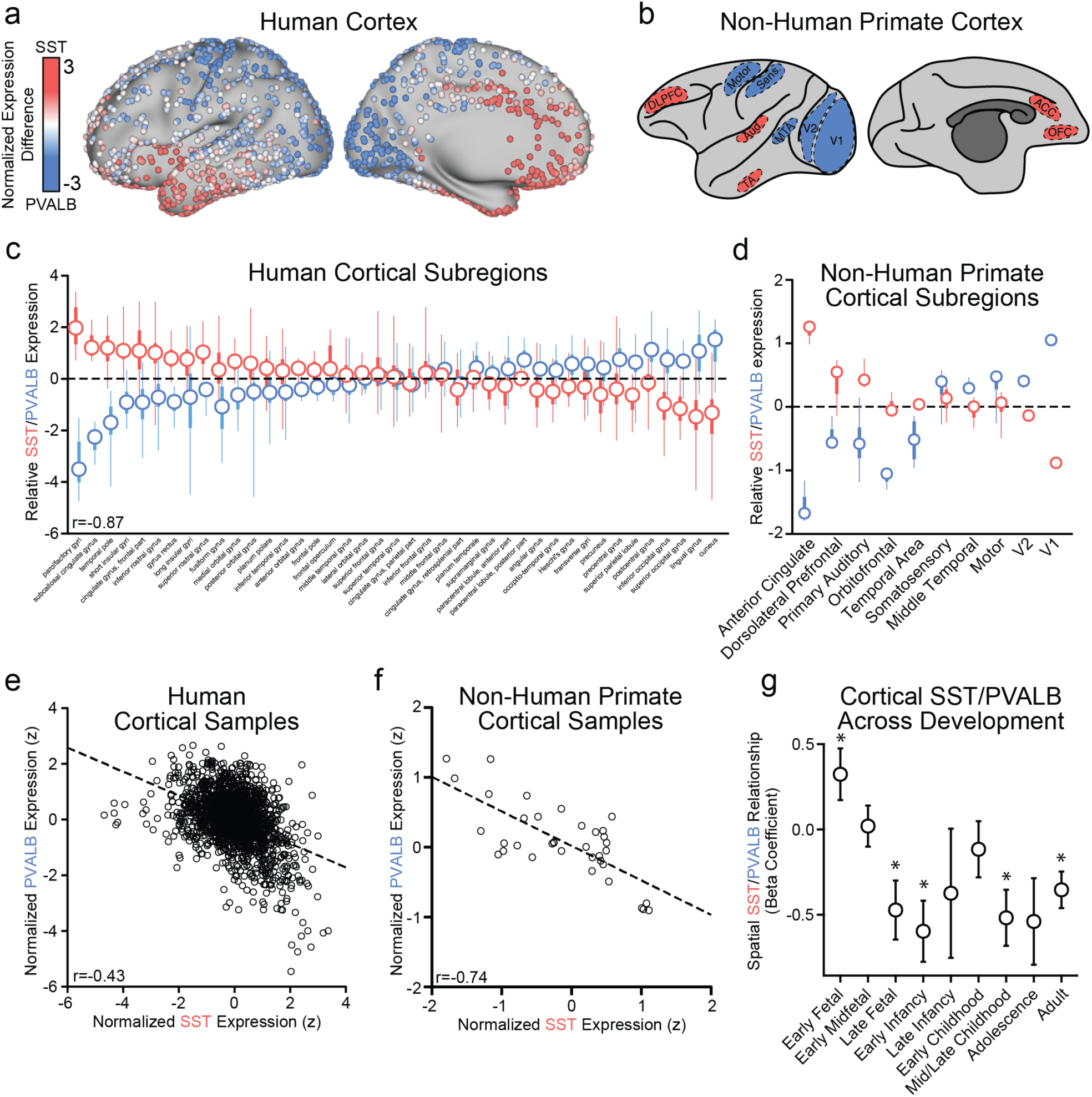
Cortical expression of *SST* and *PVALB* are negatively correlated across species and developmental stages. (a) Left-hemisphere AHBA tissue samples mapped to the human cortical surface, and (b) an illustration of non-human primate tissue sample locations, colored by relative expression of *SST* (red) and *PVALB* (blue). Normalized expression difference reflects the sample-wise subtraction of mean-normalized *PVALB* from *SST*. Relative *SST-PVALB* expression among anatomically defined groups from the (c) AHBA and (d) NIH Blueprint Non-Human Primate Atlas; circles=median, thick lines=interquartile range, thin lines=min and max values. (e) Sample-wise negative correlation of *SST* and *PVALB* in (e) human cortex (r=-0.43, p≤0.001) and (f) non-human primates (r=-0.74, p≤0.001). (g) Correlation of cortical *SST* and *PVALB* across nine developmental stages using data from the Brainspan Atlas of the Developing Human Brain. *p≤0.05, uncorrected; error bars=SE.

Suggesting that interneuron spatial gradients are a core organizational feature of primate cortex, the negative spatial relationship between *SST* and *PVALB* was evolutionarily conserved in non-human macaque primates across individual samples (*r*(34)=-0.74, p≤0.001). Given that *SST* and *PVALB* interneurons originate along a stereotyped, negatively correlated spatial gradient in embryonic ganglionic eminences^32^, we analyzed RNAseq data from the Brainspan Atlas of the Developing Human Brain to test whether *SST*/*PVALB* negative gradients emerge during developmental periods coinciding with major waves of interneuron colonization, approximately 10-25 post conception weeks(pcw)^35,36^. The negative correlation between *SST* and *PVALB* was absent in early-fetal (8-12 pcw; β=0.32, p=0.04) and early-midfetal (13-21 pcw; β=0.02, p=0.85) ages. Consistent with the hypothesis that mature interneuron distributions result from developmentally programmed migration patterns, we observed significant negative correlations between *SST*/*PVALB* across late-fetal (24-37 pcw; β=-0.47, p=0.012), early-infancy (4 months; β=-0.60, p=0.0033), mid-late childhood (8-11yrs; β=-0.52, p=0.0038), and adult (18-40yrs; β=- 0.35, p=0.0014) age, although not in late infancy (10 months; *b*=-0.37, p=0.36), early-childhood (1-4yrs; *b*=-0.11, p=0.48), or adolescence (13-15yrs; *b*=-0.54, p=0.057). These data provide developmental context as well as an external replication of the *SST/PVALB* cortical expression pattern observed in the AHBA (adult human) and NHP Atlas (adult macaque) samples.

### *SST* and *PVALB* distinguish limbic and somato/motor cortico-striato-thalamic networks

Spatial patterns of gene expression may recapitulate the architecture of functional networks across cortex^8,10,37^ and mirror functional connectivity between territories with vastly different global expression profiles (e.g., cortex and striatum^9^). We next examined whether the inverse spatial relationship between *SST* and *PVALB* is unique to cortex or preserved across subcortex (See Supplemental Information for subcortical sample information). Sample-wise expression was normalized separately for each of seven areas: striatum, thalamus, hypothalamus, globus pallidus, amygdala, hippocampus proper (i.e. CA1-CA4), and combined substantia nigra/ventral tegmentum. A cumulative negative relationship was observed between *SST* and *PVALB* (Figure 2a; r=-0.14), although a wide range of correlation values were observed (from -0.71 through 0.29). To demonstrate that the observed overall negative correlation between *SST* and *PVALB* is not obligated by global transcriptional properties, we display the distribution of averaged correlations, collapsed across cortex and the seven subcortical areas, of every gene to *SST* and to *PVALB*. Figure 2b demonstrates that *SST* is among the most negatively correlated genes to *PVALB* (bottom 0.014% of distribution), across all regions. Similarly, *PVALB* is among the most negatively correlated genes to *SST* (bottom 0.0012% of distribution). See Supplemental Figure 2 for *SST* and *PVALB* expression across subcortical subregions.

**Figure 2.**
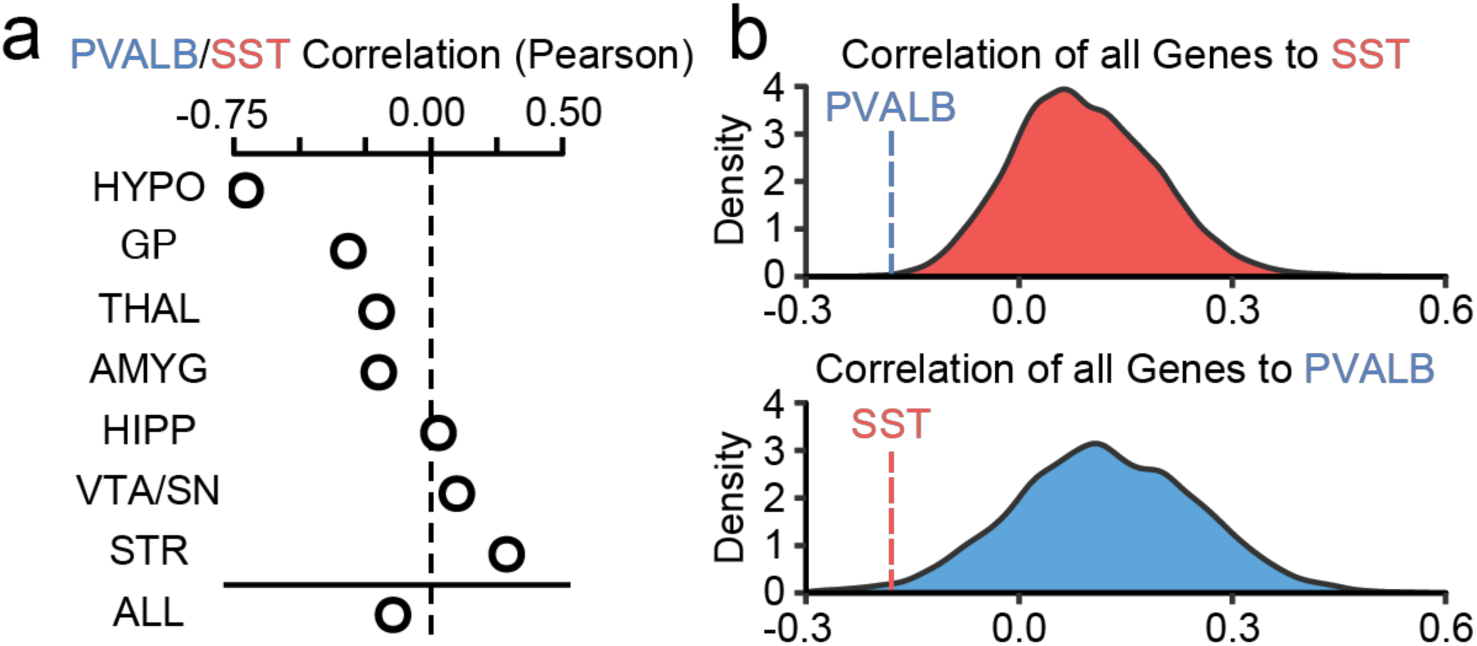
SST and PVALB are among the most negatively correlated transcripts to one another. (a) Overall, *SST* and *PVALB* are negatively correlated in subcortical regions (r=-0.14), but this relationship is variable (range -0.71–0.29). (b) The spatial correlation between *SST* and *PVALB* was averaged across cortex and the seven analyzed subcortical regions. Compared to all genes, *SST* and *PVALB* are among the most negatively correlated genes to one another. *PVALB* is among the top 0.0012% most negatively correlated genes to *SST* (top panel), and *SST* is among the top 0.014% most negatively correlated genes to PVALB (bottom panel). HYPO=hypothalamus, GP=Globus Pallidus, THAL=thalamus, AMYG=amygdala, HIPP=hippocampus, VTA/SN=ventral tegmental area/substantia nigra, STR=striatum, ALL=averaged subcortical correlation.

Although some subcortical regions display a positive *SST*-*PVALB* spatial correlation (e.g. striatum), anatomically defined regions do not always reflect functional organization^38^. For instance, the putamen contains subregions that differentially couple with default, frontoparietal, and somato/motor cortical functional networks^39^. Consequently, interneuron markers may show stereotyped patterns of expression when viewed through the lens of global functional network architecture rather than anatomy. Parallel distributed networks connect cortex, striatum, and thalamus to support complex affective, cognitive and motor behaviors^40^. To characterize relationships between these network boundaries and interneuron subtype organization, AHBA samples were aligned and analyzed according to functional parcellations of the striatum^39^ and thalamus^41^. Suggesting that interneuron subtypes differentiate large-scale functional networks, paired-sample *t*-tests revealed significantly greater expression of *SST,* relative to *PVALB,* within a distributed limbic network encompassing ventral striatum (t(15)=6.08, p=2.1e-5), mediodorsal thalamus (t(72)=2.41, p=0.018; Figure 3c), and subgenual anterior cingulate and medial prefrontal cortex (mPFC; Figure 3e). Furthermore, we established that *SST*-biased sub-regions of the thalamus and striatum (Figure 3e-f) form a distributed functional network using resting-state data from an adult community-based sample (percent female=54.47, Age=62.66 (SD 7.45), min=45, max=80) from the UK Biobank project (N=9,627 see Supplemental Figures 3 & 4 for cortical correlations). Limbic striatum and default thalamus (Figure 3c) displayed overlapping positive functional connections (*r*’s>0.05) within medial prefrontal cortex (Figure 3e), an area with strong preferential expression of *SST* (*F*(1,337)=14.09, p=0.0002; Figure 3g). This mPFC-ventral striatum-mediodorsal thalamus network broadly supports reward and affective information processing and is consistently implicated in affective disorders^42^.

**Figure 3.**
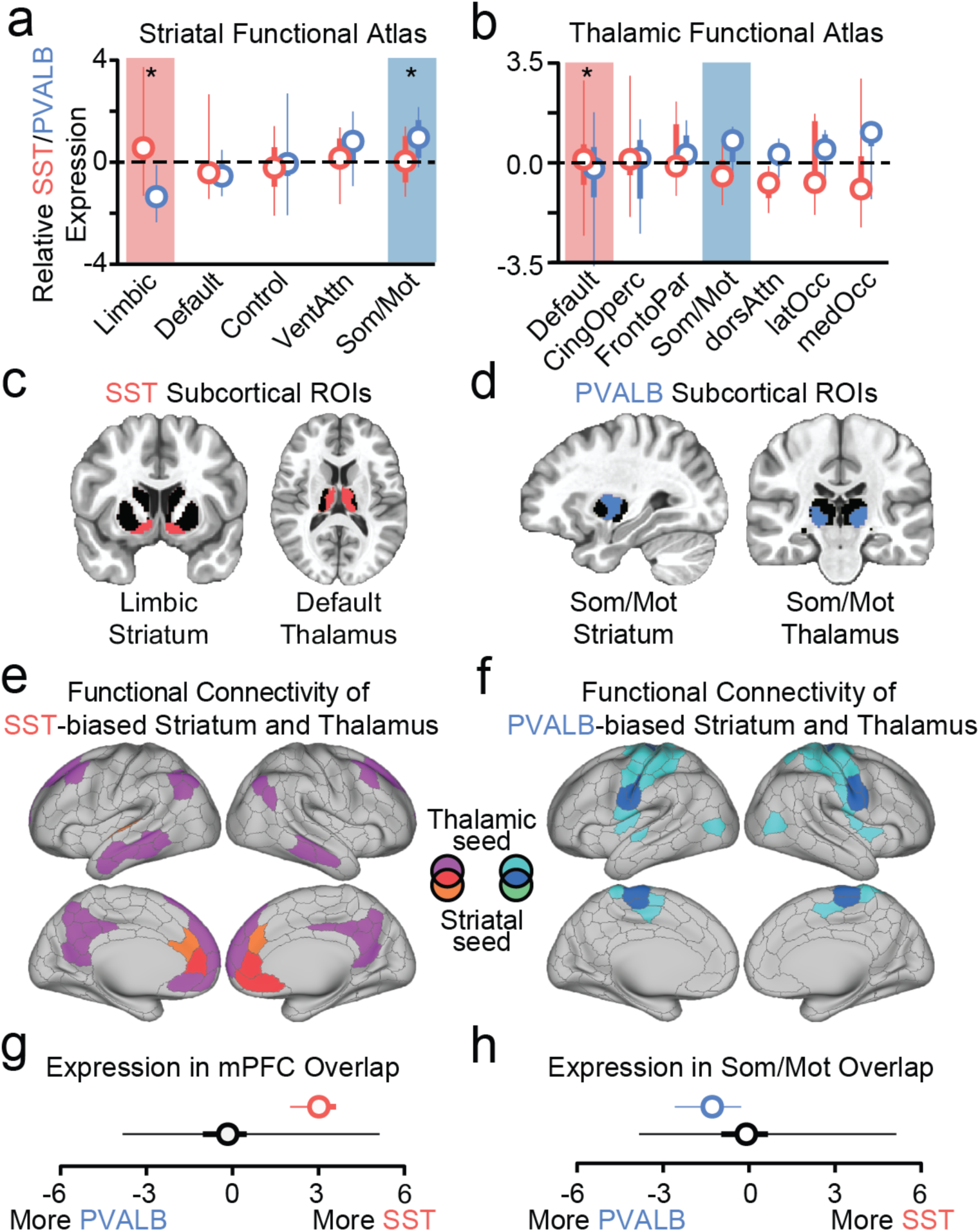
Differential *SST*/*PVALB* expression in distributed limbic and somato/motor networks. (a and b) Relative *SST* and *PVALB* expression across functionally defined striatal and thalamic subregions.(c) Relative expression of SST was highest within limbic striatum and default thalamus. (d) Relative PVALB expression was greater within somato/motor striatum and somato/motor thalamus. (e) Limbic striatum and default thalamus possess overlapping positive resting state correlations to *SST*-biased aspects of medial prefrontal cortex (mPFC; *r’s*≥0.05). Likewise, (f) somato/motor striatum and thalamus show overlapping positive correlations to *PVALB*-biased portions of somatosensory cortex (*r’s*≥0.03). (g) *SST* expression within the 3 overlapping mPFC limbic parcels is greater than all other cortical parcels. (h) *PVALB* expression within the 15 overlapping somato/motor parcels is greater than all other cortical parcels.

Among regions of a distributed somato/motor network, relative *PVALB* expression was increased within sensory and motor cortex (Figure 3f) and dorsolateral putamen (t(11)=3.47, p=0.031), but not within ventrolateral thalamus (t(5)=1.22, p=0.28; Figure 3d), which may be due to particularly sparse sampling in this region (n=5). A visual medial occipital area of the thalamus also displayed preferential expression of *PVALB* (t(14)=2.74, p=0.016; Figure 3b), consistent with the proposal that *PVALB/SST* ratios are higher in distributed whole-brain networks that process visual and sensorimotor information^13^. Supporting this distinction, both somato/motor striatum and thalamus (Figure 3d) were positively functionally coupled (*r*’s>0.03) to motor and sensory cortex (Figure 3f), which show a *PVALB* expression bias (*F*(1,337)=6.86, p=0.009; Figure 3h).

Suggesting the *SST/PVALB* dissociation also extends to the midbrain, relative expression of *SST* (M=1.15 SD=0.52) was greater than that of *PVALB* (M=-0.24 SD=0.71) among ventral tegmental area (VTA) samples (t(12)=-7.17, p=1.1e-5), whereas *PVALB* (M=0.69 SD=0.88) was greater than *SST* (M=- 0.13 SD=0.90) in the substantia nigra reticulata (STNr; t(23)=2.98, p=0.006; Supplemental Figure 2). The VTA is densely interconnected to other *SST*-biased regions, including the nucleus accumbens (NAcc), anterior cingulate cortex, and mediodorsal thalamus^42^, whereas functional neuroimaging and tract-tracing work suggests the substantia nigra pars reticulata (SNr) preferentially functionally couples to motor areas and is reciprocally connected to sensorimotor striatum^40,43,44^. Together, these data suggest that *SST* expression is greater within distributed limbic and affect-related regions, whereas *PVALB* expression is elevated within a distributed sensorimotor processing network.

### *SST*/*PVALB* ratios co-vary with resting-state signal amplitude across cortex

Computational work in rodents posits that the ratio of SST to PVALB interneurons contributes to regional differences in function and hierarchical organization across cortex^13^. Sensory and association cortices display hierarchically organized timescales of spiking activity that progress from shorter to longer, respectively^17,45^. This aspect of functional organization may be indexed by variability in the resting-state BOLD signal. Accordingly, we examined whether the ratio of cortical *SST*/*PVALB* expression relates to an *in-vivo* measurement of cortical signal variability, resting-state functional amplitude (RSFA)^46^. Voxel-wise RSFA was calculated using the UK Biobank sample (n=9,627) and averaged across the 400 parcel functional atlas of Schaefer and colleagues^47^.

We first established the heritability of RSFA. Between-subjects hierarchical clustering was used to identify cortical territories with similar patterns of signal amplitude across individuals (Figure 4a), corresponding to limbic (light beige), cingulo-opercular (teal), temporo-parietal (orange), prefrontal (red), somato/motor (blue), and visual (purple) clusters. Consistent with recent work^48^, this data-driven dimensionality reduction broadly categorized association and unimodal aspects of cortex. Suggesting that individual differences in RSFA can be explained by genetic variation in the general population, a significant proportion of between-subject variation in cluster-wise RSFA was found to be due to common genetic factors [h^2^_snp_: limbic=0.27 (SE 0.04), cingulo-opercular=0.22 (SE 0.04), temporo-parietal=0.28 (SE 0.04), prefrontal=0.31 (SE 0.04), somato/motor=0.21 (SE 0.04), visual=0.06 (SE 0.04)]^49^. See Supplemental Figure 5 for parcel-wise estimates of RSFA heritability.

**Figure 4.**
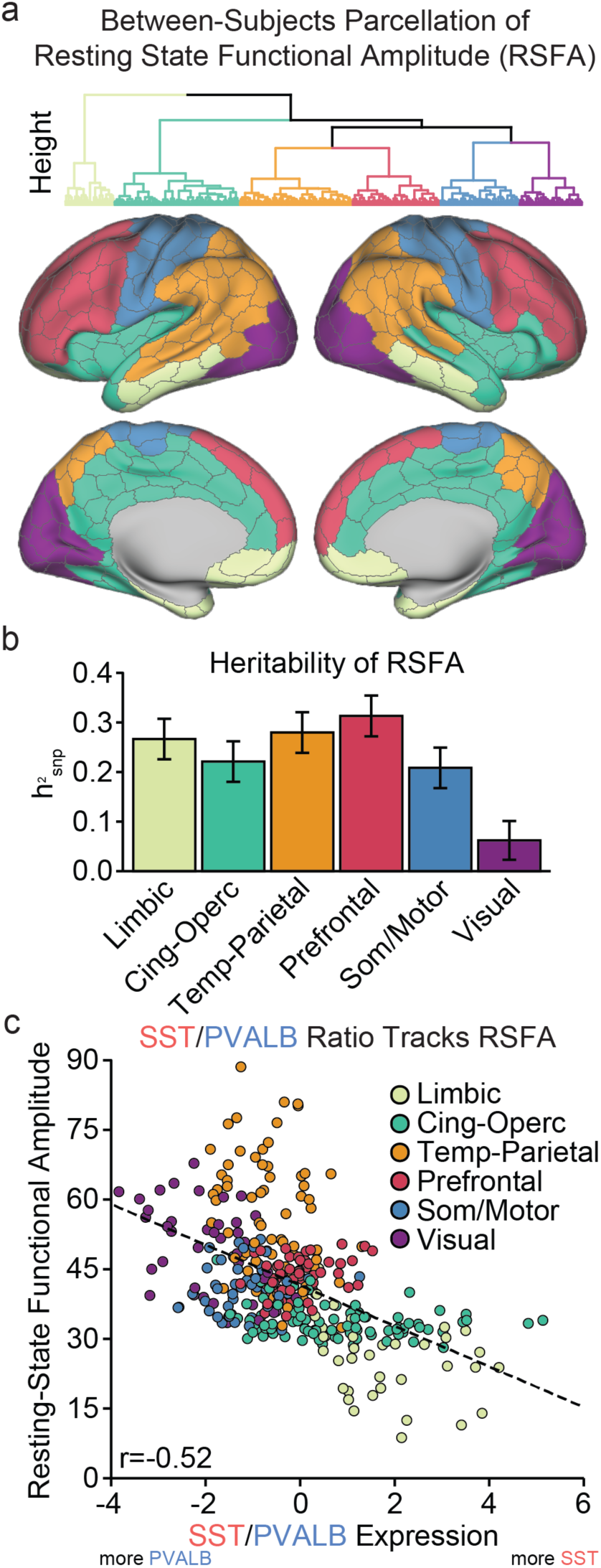
*SST/PVALB* ratio tracks inter-regional differences in cortical brain function. (a) Mean RSFA was calculated for each of 400 volumetric cortical parcels from the Schaefer parcellation^47^.Between-subjects hierarchical clustering of residualized RSFA values revealed 7-clusters of parcels with similar amplitude signatures; Light beige=limbic, teal=cingulo-opercular, orange=temporal-parietal, red=prefrontal, blue=somato/motor, and purple=visual. (b) Overall, a significant proportion of variability of RSFA was explained by common genetic variation (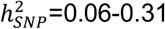; error bars=standard error) (c) Parcel-wise relative expression of *SST*/*PVALB* is negatively correlated to RSFA (*r*(337)=-0.52, p≤2.2e-16).

**Figure 5.**
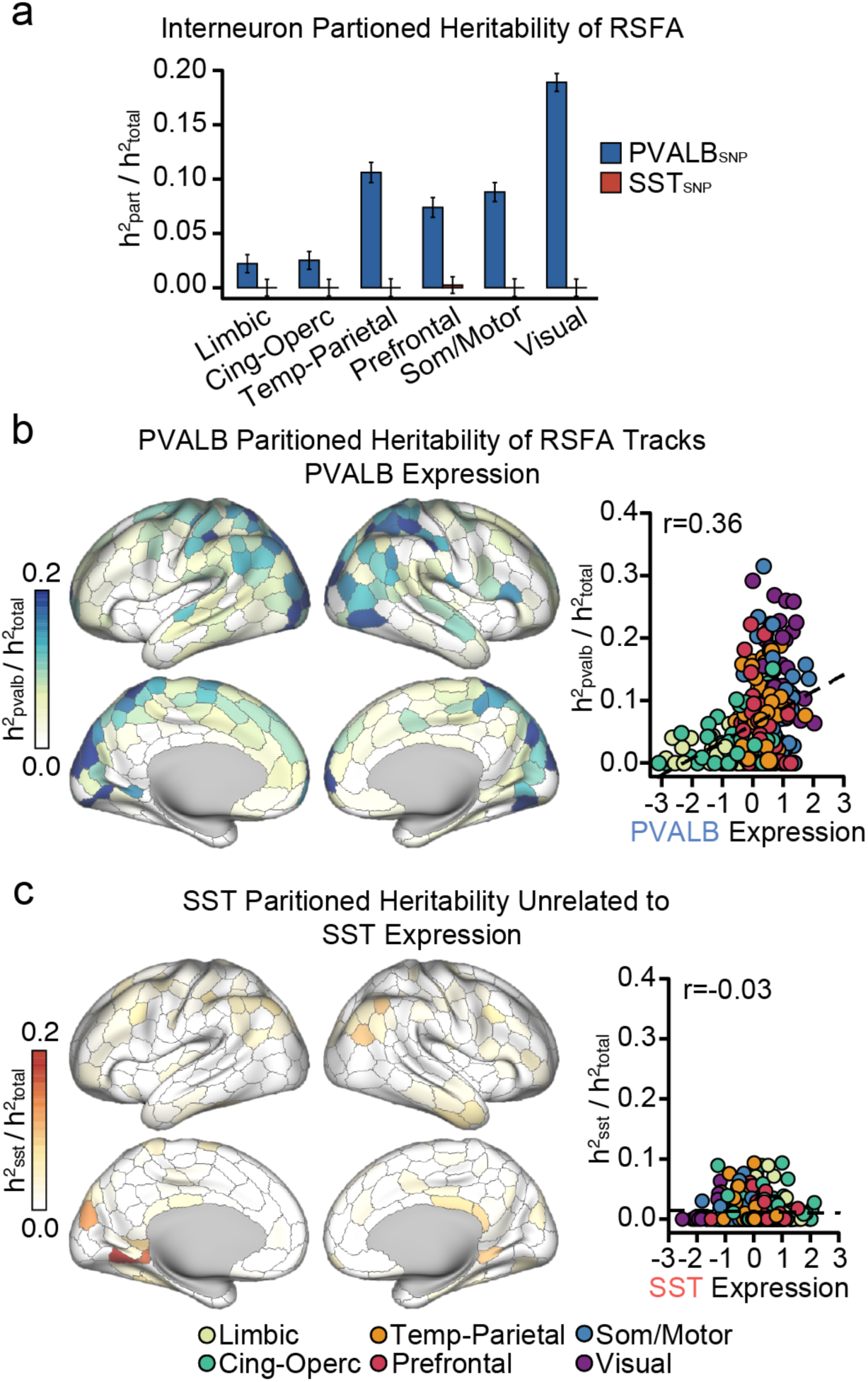
*PVALB* genes underlie spatially variable patterns of heritable brain function. (a) Across the six data-defined RSFA clusters, the 500 gene PVALB_SNP_ set accounted for a significant proportion of heritable variance in cingulo-opercular, prefrontal, and temporo-parietal areas. PVALB_SNP_ enrichment was observed within tempero-parietal, prefrontal, som/motor, and visual clusters. (b) Parcel-wise partitioned heritability tracks sub-type specific gene expression for PVALB_SNP_ (*r*(326)=0.36, p=1.78e-11), (c) but not the SST_SNP_ (*r*(326)=-0.03, p=0.54), partitions. Error bars=SE.

Expression data from the AHBA were used to test whether interneuron ratios track the spatial layout of RSFA signal variability across the cortical sheet. Earlier work has documented a correlation of interneurons marker expression with fractional Amplitude of Low-Frequency Fluctuations (fALFF)^50^, a metric closely tied to RSFA, within a circumscribed set of cortical areas. Across the whole-brain Schaefer cortical parcellation, *SST*/*PVALB* ratio was negatively correlated with resting-state signal amplitude (*r*(337)=-0.52, p≤2.2e-16; Figure 4c. Parcels with higher relative expression of *SST* had lower RSFA (e.g. limbic parcels). Conversely, clusters with higher relative *PVALB* had higher RSFA (e.g., visual, parietal). Across individual interneuron markers, we observed a positive correlation to parcel-wise RSFA and expression of *PVALB* (r=0.48, p≤2.2e-16), and a negative correlation to *SST* (r=-0.44, p≤2.2e-16).

### Polygenic variation among *PVALB*-correlated genes underlies cortical brain function

Genome-Wide Association Studies (GWAS) demonstrate that the genetic bases of many complex traits are due to the cumulative weight of genetic variants spread across the entire genome, each with a subtle effect^51^. Although brain phenotypes such as resting-state functional amplitude likely display such a polygenic architecture^52^, phenotype-relevant polymorphisms can cluster in genes expressed within relevant tissue and cell types^53^. Given that cortical resting-state functional amplitude tracks the topography of interneuron ratios, we next tested whether single-nucleotide polymorphisms (SNPs) underlying the heritable variance in brain activity (i.e. RSFA) are enriched within genes linked to *PVALB* and *SST*. The observation that RSFA-related SNPs are enriched within interneuron-related genes would yield insight into the molecular basis of the resting-state BOLD fluctuations.

Interneuron-correlated gene sets were nominated using a guilt-by-association logic. That is, genes that were spatially correlated to interneuron markers (i.e. *SST, PVALB*) were assumed to relate to each interneuron subtype. Using cortical AHBA data, genes were rank-ordered based on their spatial correlation to each interneuron marker and the top 500 most-correlated genes were selected. *PVALB* and *SST* gene sets were non-overlapping. Interneuron-related SNP lists were generated for each gene set by identifying variants within ±5000 base pairs from transcription start and stop site of each gene. eQTL variants for each gene set were included, defined using cortical data from the CommonMind consortium^54^ and NIH GTEx^55^. We denote the SNP lists for each interneuron gene set as PVALB_SNP_ and SST_SNP_ (see Supplemental Data). Genetic relatedness matrices were calculated for the UKB sample using each SNP set, and heritability was estimated using GCTA-REML simultaneously across three partitions: PVALB_SNP_, SST_SNP_, and a partition containing all remaining genotyped variants^49^.

Indicating that the genetic basis of RSFA, a measure of *in vivo* brain activity, is determined in part by genes linked to *PVALB* interneurons, the PVALB_SNP_ set accounted for a significant proportion of heritable variance of the temporo-parietal (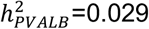, SE=0.0092, q=0.00047) prefrontal (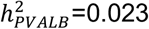, SE=0.0091, q=0.016), and somato/motor (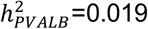, SE=0.0087, q=0.034) RSFA clusters, but not the limbic (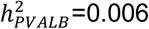, SE=0.008, q=0.25), cingulo-opercular (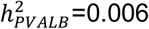, SE=0.0082, q=0.25), or visual (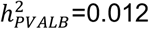, SE=0.008, q=0.11) clusters. Conversely, the SST_SNP_ set did not explain a significant proportion of heritable variance across any partition (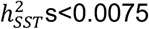, ps>0.46). A key question is whether the genetic variance explained by the PVALB_SNP_ set is greater than what is expected given the number of SNPs examined, which would indicate the outsized, or enriched, role of these genetic variants in RSFA. Enrichment was calculated as the proportion of heritability explained by the partition, divided by the fraction of SNPs in that partition, where a value greater than 1 denotes enrichment. We observed fold enrichment greater than 1 for visual (enrich=6.49 SE=0.28), motor (enrich=3.02 SE=0.30), temporo-parietal (enrich=3.64 SE=0.32), prefrontal (enrich=2.53 SE=0.31) clusters, but not limbic (enrich=0.76 SE=0.29) or cingulo-opercular (enrich=0.86 SE=0.28). The PVALB_SNP_ list (N=9,819 variants) constituted 2.9% of total analyzable genotyped SNPs (N=337,356 variants), but accounted for 2.2-18.9% (M=8.4 SD=6.1) of total genetic variance across each of the RSFA clusters (Figure 5a). The SST_SNP_ partition (2.6% of available variants) did not explain a significant proportion of genetic variance for any RSFA cluster.

An important unanswered question is whether the genetic determinants of RSFA are uniform across cortex, or whether they vary according to underlying cytoarchitecture. We next tested whether the PVALB_SNP_ and SST_SNP_ partitions explain a greater percentage of heritable RSFA variance in regions where the respective marker is expressed most. Partitioned heritability analyses were performed for each of the 400 Schaefer cortical parcels. Across all parcels with available AHBA expression data, normalized genetic variance explained by the PVALB_SNP_ partition was positively correlated to *PVALB* expression (Figure 5b; *r*(326)=0.36, p=1.78e-11), corresponding to visual, superior temporal, and parietal areas of cortex (Figure 5b). Across all genes, *PVALB* was among the top 64 transcripts (top 0.003% of 20,738 transcripts) showing a positive spatial correlation to the PVALB_SNP_ partition map (i.e. Figure 5b), indicating that this positive relationship is not obligated by global statistical properties. Conversely, partitioned PVALB_SNP_ heritability was negatively correlated to *SST* expression (*r*(326)=-0.36, p=2.85e-11). There was not a significant parcel-wise relationship between SST_SNP_ partitioned heritability and SST gene expression (*r*(326)=-0.03, p=0.54). Together, these findings indicate that the molecular genetic basis of resting-state functional amplitude is spatially heterogeneous, demonstrating a particularly important role of genes co-expressed with *PVALB*.

### The association between genetic risk for schizophrenia and brain function follows the spatial profile of *PVALB* expression

Understanding the molecular genetic underpinnings of brain function is pressing given the need for empirically informed treatment targets for heritable, brain-based, psychiatric illnesses like schizophrenia^56^. Convergent evidence from animal models and post-mortem tissue analyses suggests that interneuron dysfunction as a core pathophysiological feature of schizophrenia^57^. To determine whether interneuron-related genetic variation is tied to disease liability, we tested whether polygenic risk for schizophrenia^58^ is greater among PVALB_SNP_ and SST_SNP_ variants, relative to the rest of the genome. Using a partitioned MAGMA analysis^59^, we divided rank-ordered *PVALB,* and *SST* gene lists into bins of 500. Using MAGMA, we observed significant enrichment of schizophrenia polygenic risk for the top *PVALB* gene set (beta=0.083, p=0.038), but not the top *SST* (beta=-0.01, p=0.61). Suggesting that polygenic schizophrenia risk is greater among interneuron-related genes, we examined all gene bins examined and found that the enrichment of schizophrenia genetic risk decreased as gene bins became less spatially correlated with *PVALB* (*r*(18)=-0.65, p=0.0017), but not *SST* (*r*(18)=-0.39, p=0.09; Figure 6a).

**Figure 6.**
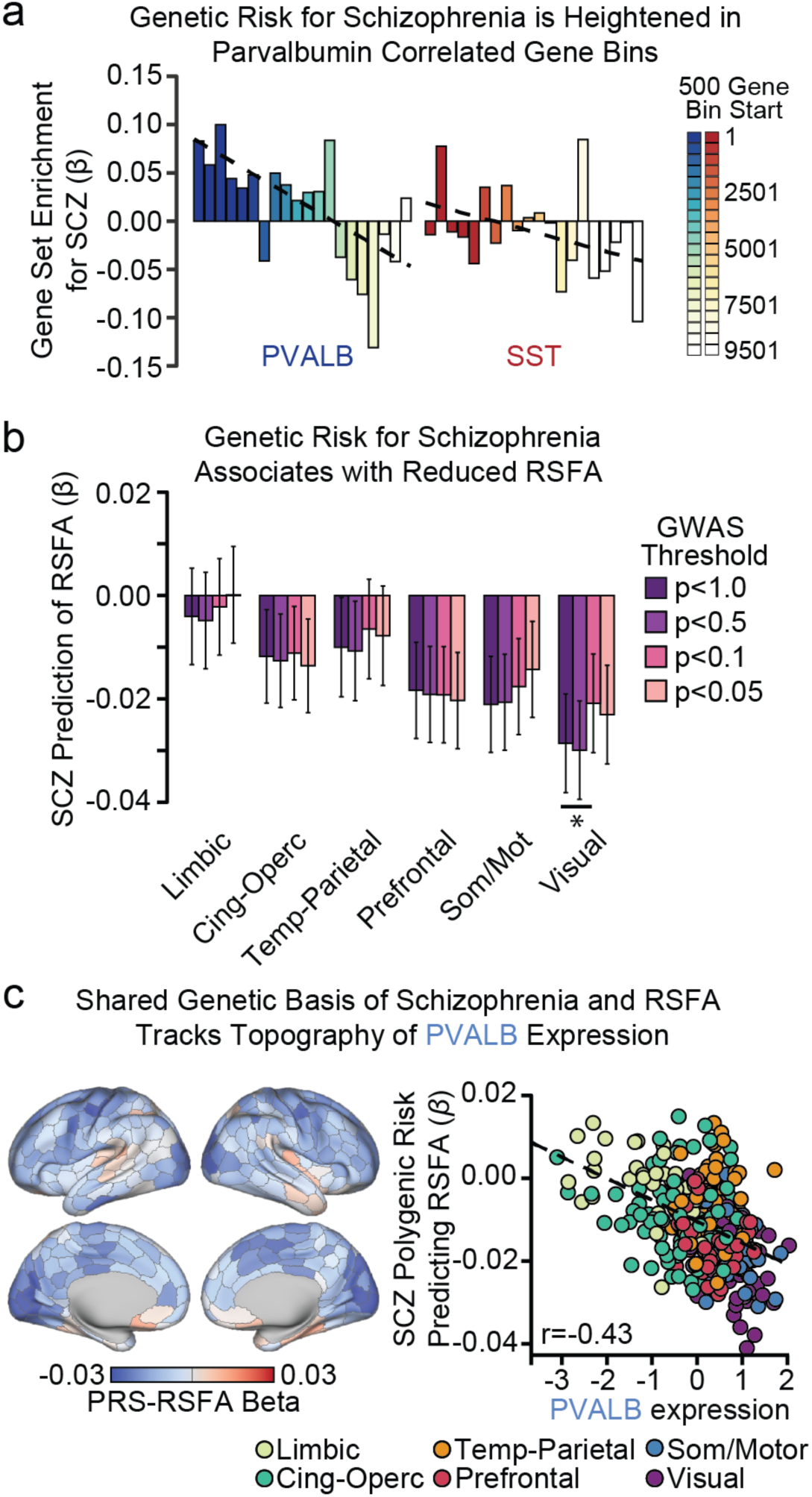
Schizophrenia polygenic risk predicts brain function and tracks *PVALB* expression. (a) Genes were rank-ordered by cortical spatial correlation to *SST* and *PVALB,* then divided into 500 gene bins. MAGMA competitive gene set analysis revealed enrichment of polygenic risk for schizophrenia in the top *PVALB* (p=0.032), but not the top *SST* (p=0.62) set. Enrichment decreased across ordered bins for *PVALB* (r=-0.65, p=0.0017) and *SST* (r=-0.39, p=0.09). (b) Schizophrenia polygenic risk negatively predicts RSFA within the visual (q=0.016) cluster, as well as somato/motor (q=0.070) and prefrontal (q=0.097) clusters at trend-levels. (f) Parcel-wise prediction of RSFA by the schizophrenia PRS significantly negatively correlated with cortical expression of *PVALB* (r=-0.43, p=2.2e-16). SCZ=schizophrenia; PRS=Polygenic Risk Score; RSFA=Resting State Functional Amplitude. *=q≤0.05.

To test whether polygenic risk for schizophrenia influences cortical RSFA, we calculated a schizophrenia polygenic risk score (SCZ-PRS)^60^ using genotyped variants from individuals in the UK Biobank imaging sample. Across the data-derived RSFA clusters, SCZ-PRS negatively predicted RSFA in the visual cluster (Benjamini-Hochberg corrected q=0.016; Figure 6b; GWAS threshold p<1.0), as well as somato/motor (q=0.070) and prefrontal (q=0.097) clusters at trend-levels. Consistent with the hypothesized link between PVALB interneurons and psychotic illness, the relationship between RSFA and polygenic schizophrenia risk was significantly negatively correlated to the topography of *PVALB* expression across cortex (*r*(337)=-0.43, p≤2.2e-16; Figure 6c). That is, regions where SCZ-PRS most negatively predicted RSFA corresponded to areas with the greatest *PVALB* expression (e.g. motor and visual parcels). This relationship remained significant after controlling for the overall SNP-wise heritability of each parcel (β=-.38, *t*(336)=-7.48, *p*=6.73e-13), indicating that the effect is independent of parcel-wise explainable genetic variance. Comparing the RSFA-schizophrenia polygenic risk map to all genes, *PVALB* was the among the top 0.0034% negatively correlated expression profiles (72 out of 20,738), showing that this relationship is not obligated by globally negative relationships between gene expression and schizophrenia risk RSFA effects. Ontological enrichment analysis further revealed that the top 500 genes correlated with *PVALB* in the AHBA data contained genes associated to schizophrenia and bipolar disorder, neuronal signaling, and gated channel activity (Table 1). Together, these data suggest that schizophrenia-related genetic variants cluster within cell types, particularly parvalbumin interneurons, leading to differential functional disruption across cortex.

**Table 1.** Enrichment terms for interneuron-correlated genes. Ontological enrichment analyses were conducted with ToppGene on the 500 genes used to generate the PVALB_SNP_ and SST_SNP_ lists.

|  | Category | ID | Name | p | q FDR-BH | Hits | Genes in GO |
| --- | --- | --- | --- | --- | --- | --- | --- |
| SST | GO: BP | GO:0099536 | synaptic signaling | 1.08e-5 | 2.16e-2 | 37 | 687 |
|  | GO: BP | GO:0099537 | trans-synaptic signaling | 1.95e-5 | 2.16e-2 | 36 | 678 |
|  | GO: CC | GO:0097458 | neuron part | 3.35e-6 | 6.31e-4 | 68 | 1545 |
|  | GO: CC | GO:0045202 | synapse | 3.3e-4 | 1.69e-2 | 39 | 870 |
| PVALB | GO: MF | GO:0005261 | cation channel activity | 3.33e-12 | 1.74e-9 | 33 | 306 |
|  | GO: MF | GO:0005249 | voltage-gated potassium channel activity | 1.33e-10 | 1.69e-8 | 17 | 91 |
|  | GO: BP | GO:0071805 | potassium ion transmembrane transport | 2.75e-10 | 6.61e-7 | 23 | 181 |
|  | GO: BP | GO:0098655 | cation transmembrane transport | 9.85e-9 | 7.88e-6 | 47 | 738 |
|  | GO: CC | GO:0034703 | cation channel complex | 1.04e-10 | 5.07e-8 | 23 | 176 |
|  | GO: CC | GO:0034702 | ion channel complex | 3.03e-10 | 5.07e-8 | 29 | 291 |
|  | Disease | C0036341 | Schizophrenia | 1.93e-7 | 7.27e-4 | 70 | 1561 |

## Discussion

Integrating genetic, transcriptional, and neuroimaging data, we demonstrate that spatial distributions of interneurons are stereotyped across species and development, align to the topographic distribution of functional brain networks, and underlie a substantial portion of the heritable aspects of resting-state functional amplitude, a measure of *in vivo* brain activity. Somatostatin- and parvalbumin-interneuron markers were negatively spatially correlated across cortex, a relationship that was robust in early developmental periods in humans and evolutionarily conserved in non-human primates (Figure 1). Stereotyped patterns of *SST* and *PVALB* expression were observed in subcortex (Figure 2), with *SST* and *PVALB* differentially expressed within distinct limbic and somato/motor functional networks linking cortex, striatum, and thalamus (Figure 3), respectively. Computational models theorize that interneuron ratios underlie regional differences in cortical brain function^13^. Providing empirical support for this hypothesis, regional differences in *SST/PVALB* expression in post-mortem brain tissue align with spatial variability in resting-state functional amplitude in the general population (Figure 4). Suggesting the functional relevance of this spatial relationship, genetic polymorphisms linked to PVALB interneurons accounted for an enriched proportion of heritable variance underling cortical signal amplitude (Figure 5). Critically, the amount of variance explained by *PVALB* SNPs positively tracked spatial expression of *PVALB*, suggesting that common genetic polymorphisms influence brain function in a cell-type specific and regionally variable manner. Implicating genetic differences among interneurons in schizophrenia, schizophrenia-related polygenic risk was enriched among genes co-expressed with interneurons, and predicted reduced resting-state functional amplitude across cortex in a manner that tracked the spatial landscape of *PVALB* gene expression (Figure 6).

Adaptive functioning depends on the brain’s capacity to integrate information across timescales. Higher-order cognition often requires information accumulation over time, whereas sensorimotor processing entails rapid adaption to changing external stimuli^18,47,65,66^. These informational demands are met, in part, through the hierarchical organization of anatomic and functional connections in cortex, as well cytoarchitectural gradients that underlie regional specialization^12,61^. Our data indicate that interneuron ratios, as indexed by *SST* and *PVALB* expression, are an important feature underlying regional differences in brain function (Figure 4). Due to unique electrical and synaptic properties of somatostatin and parvalbumin interneurons, relative shifts in their density can alter the balance of inhibitory control^13^. SST interneurons synapse onto dendrites of pyramidal neurons to gate incoming cortical signals, whereas PVALB interneurons provide perisomatic inhibition that is well-suited for feedback inhibition and output regulation^2^. Computational models suggest that increased dendritic (i.e. SST) over perisomatic (i.e. PVALB) inhibition results in more robust filtering of distracting information, allowing for greater recurrent excitation in association cortex for complex tasks requiring integration of information over time^20^. Conversely, sensorimotor regions may benefit from fast responses and lower recurrent excitation to adapt to rapidly changing inputs^17^, which could be facilitated by direct inhibitory signals from parvalbumin-expression interneurons.

Our analyses provide molecular genetic support for a relationship between parvalbumin interneurons and the hemodynamic BOLD signal. A wealth of evidence indicates that BOLD signal most tightly couples to gamma oscillations (30-80 Hz) relative to other frequency domains^21-25^. Individual differences of GABA in visual cortex predict both gamma oscillations and BOLD amplitude^62^, a relationship that animal work suggests is primarily driven by parvalbumin interneurons^19^. Here, we provide initial evidence in humans for the preferential influence of parvalbumin interneurons on fMRI signal. For instance, polygenic variation among parvalbumin correlated genes explained upwards of 18% of the heritable variance in RSFA in visual cortex.

Schizophrenia is among the most heritable forms of psychiatric illness (*h*^2;^=81%)^63^, underscoring the pressing need to map polygenic variation to illness-related brain phenotypes and associated risk factors. Converging lines of evidence point to GABAergic abnormalities as a cardinal feature of the disorder^64^, highlighting a particular role of parvalbumin interneurons^26^. Patients with schizophrenia exhibit reduced levels of GAD67, an enzymatic precursor of GABA^65^, and are characterized by parvalbumin interneurons with atypical perineuronal nets^66^, dysregulated mitochondrial gene transcription^67^, and reduced potassium signaling channels^68^ relative to healthy populations. These abnormalities are thought to underlie disrupted gamma-band oscillations and working memory deficits which are a hallmark of the disorder^64^. Linking these observations, we demonstrate here that polygenic risk for schizophrenia is increased among genes that are spatially correlated to *PVALB* (Figure 6a), expanding upon cell transcriptomic work implicating cortical interneurons as an illness marker^53^. Consistent with a relationship between schizophrenia-linked genetic vulnerability and brain function, we document a negative association between individual polygenic schizophrenia risk and resting-state functional amplitude in a large population-based sample (Figure 6b). Importantly, the topography of these effects follows spatial profile of *PVALB* expression across cortex (Figure 6c), highlighting the potential role of parvalbumin interneurons in mediating brain-based intermediate phenotypes associated with illness risk.

Disruption of excitatory/inhibitory balance is thought to reflect a cross-diagnostic marker of psychiatric illness^69^. For instance, decreased expression of parvalbumin cell markers is evident in both schizophrenia and bipolar disorder^70^, while major depressive disorder (MDD) is marked by selective reductions in somatostatin interneurons^71^. Delineating the region-specific functional roles of cortical interneuron subtypes will provide biological insight into cross-diagnostic patterns of both behavior and brain function. With regard to depressed mood and negative affect, modulation of cortical somatostatin interneurons can causally influence anxiety- and depressive-like behavioral phenotypes in rodents^7,72^. In line with this observation, we observe the greatest expression of somatostatin within a distributed limbic network linking mPFC, NAcc, and mediodorsal thalamus (Figure 3) that processes reward and affective information^42^. Somatostatin-biased cortical regions (ACC, mPFC, and insula) also correspond to areas where cortical thinning has been observed in patients with MDD^73,74^ and individuals reporting elevated negative affect^75^. These converging lines of evidence support the hypothesized role of somatostatin neurons in mood-related psychiatric symptoms^71^, which should be explored in future work on the molecular and neural underpinnings of affective illness.

The present findings should be interpreted in light of several limitations. First, we use single molecular markers to infer the relative presence of SST and PVALB interneurons, which are not sensitive to morphological and physiological differences among interneuron subgroups^2^. More nuanced inference of cellular spatial distributions should be conducted as single-cell transcriptomic atlases are developed in humans. Further, we employ a “guilt-by-association” logic to nominate interneuron related gene sets. While we cannot conclude that genes within each identified interneuron group directly influence interneuron function, similar correlation-based nomination approaches have been shown to correspond well with *a priori* defined gene groups^76^. The examination of enrichment terms (Table 1; Supplemental Information) allows for more precise understanding of the biological processes contributing to our results. Lastly, the *in vivo* imaging and genetic analyses focus on an aging population of White/non-Latino individuals. As genetic effects can vary across ethnic and demographic subgroups^77,78^, the stability of the results reported here should be examined across diverse populations.

Inherited genetic variation shapes brain function within and across individuals^79,80^. There is pressing need to identify specific molecular genetic mechanisms of human brain function to expand our biological understanding of cognition, behavior, and associated risk for psychiatric illness. Analyses of spatially-dense, whole-genome, expression atlases increasingly reveal transcriptional correlates of brain function^50^, structure^12,87-89^, functional connectivity^8-10^, and psychiatric illness^81^. With the emergence of large-scale imaging genetic data^27^, it is now possible to bridge structural genetic, transcriptional, and large-scale neuroimaging brain phenotypes. Here, we leverage these data to show that interneuron marker distributions correlate with cortical signal amplitude, align to distributed functional networks, underlie regional differences in heritable brain function, and associate with genetic risk for schizophrenia in the general population.

## Methods

### Allen Human Brain Atlas

Publicly available human gene expression data from six postmortem donors (1 female), aged 24– 57 years of age (42.5±13.38) were obtained from the Allen Institute^33^. Data reflect the microarray normalization pipeline implemented in March 2013 (http://human.brain-map.org) and analyses were conducted according to the guidelines of the Yale University Human Subjects Committee. Microarray probes from eight overarching ontological categories were selected: cortex, dorsal thalamus, striatum, globus pallidus, hypothalamus, hippocampus proper (i.e. CA1-CA4), amygdala, and the combined substantia nigra and ventral tegmentum (see Supplemental Information). For genes with duplicate probes, the *collapseRows* function^82^ was used in R to select the probe with the highest mean expression (connectivityBasedCollapsing=FALSE), resulting in 20,738 unique mRNA probes. ComBat was used to normalize expression across donors before combining data from each brain^83^.

Individual cortical tissue samples were mapped to each AHBA donor’s Freesurfer derived cortical surfaces, downloaded from Romero-Garcia and colleagues^84^. Native space midthickness surfaces were transformed to a common fsLR32k group space while maintaining the native cortical geometry of each individual donor. The native voxel coordinate of each tissue sample was mapped to the closest surface vertex using tools from the HCP workbench^85^. Microarray expression of each gene was mean- and variance-normalized (i.e., divided by standard deviation) separately for each of the eight analyzed regions, revealing relative expression differences within cortical and subcortical territories. For region-wise expression analyses (e.g. Figure 1c), ontological categories from the AHBA were used to calculate the median, min-max, and interquartile range of relative expression in each region. Detailed information about the analyzed regions is provided in the Supplemental Information. Cortical data visualization was carried out using *wb_view* from the HCP workbench^85^. The MNI locations of striatal and thalamic samples were cross-referenced to functional atlases of Choi and colleagues^39^ and Hwang and colleagues^41^. With AFNI, a single voxel (1 mm^3^) region of interest (ROI) was generated at the MNI location of each sample. A functional network label was assigned if the ROI fell within a volumetric parcel. If the sample did not overlap with the functional atlas, the associated ROI was expanded to 2 mm^3^ and the network with the most overlapping voxels in the ROI was assigned. If the expanded 2 mm^3^ ROI did not overlap, the process was repeated using a 3 mm^3^ ROI. A sample was omitted from analysis if the 3 mm^3^ ROI did not overlap with the associated functional atlas. Functional sub-regions with 3 or fewer samples were excluded from analyses.

### UKB imaging processing

Minimally preprocessed resting-state fMRI data from the UK Biobank were analyzed, reflecting the following preprocessing steps: motion correction with MCFLIRT^86^, grand-mean intensity normalization, highpass temporal filtering, fieldmap unwarping, and gradient distortion correction. Noise terms were identified and removed using FSL ICA+FIX^87^. Full information on the UKB preprocessing is published^27^. Additional processing was conducted in AFNI^88^ and consisted of 3dDespike, resampling to MNI152 space using the UKB generated linear and nonlinear transforms, FWHM blur of 4.0mm, regression of CSF,WM, and global resting state signals, and first and second order trend removal. Voxel-wise RSFA maps were generated with 3dRSFC and then averaged within each of the approximately symmetrical 400 volumetric parcels from the 7-Network parcellation of Schaefer and colleagues^47^. Due to signal blurring between lateral striatum and insular cortex, resting-state analyses reflect an additional local white-matter regression against gray matter using AFNI anaticor. Imaging analyses were conducted in volume, but visualized on the cortical surface. Resting-state functional connectivity between striatum, thalamus, and cortex was estimated using AFNI’s 3dNetCorr, which calculated the Fisher-Z transformed correlation values of timeseries across the Choi 7-region striatal atlas^39^, the Hwang 9-region thalamic atlas^41^, and the Schaefer 400-region cortical atlas^47^.

A total of 13,236 UKB subjects were processed through the imaging pipeline. Subjects with mean run-wise frame-to-frame head motion greater than 0.20mm, and inverted rsfMRI SNR greater than 3 standard deviations above the mean were removed. After filtering for White/Non-Latino subjects with usable genetic data, cryptic relatedness <0.025, and conducting row-wise deletion for the variables age, sex, height, weight, BMI, combined gray/white matter volume, combined ventricular/CSF volume, diastolic and systolic blood pressure, run-wise rsfMRI motion, rsfMRI inverse SNR, T1 inverse SNR, and UK Biobank assessment center, 9,627 subjects remained for analyses (percent female=54.47, mean age=63.33 SD=7.45, min/max age=45-80).

### UKB genetics

UK Biobank genotype data was filtered to include only White/Non-Latino subjects with imaging data passing the quality control thresholds described above. Plink v2.00 was used to remove samples with missingness >0.10, SNPs with minor-allele frequency <0.05, Hardy-Weinberg equilibrium <1x10^-6^, and call rate <0.02, resulting in 337,356 autosomal variants^89^. GCTA software was used to calculate a genetic relatedness matrix to remove individuals with cryptic relatedness more than 0.025, leaving N=9,627 subjects for analysis^49^. Ten genetic principal components were then calculated for use as covariates in polygenic risk score and heritability analyses.

### RSFA between-subjects clustering and heritability

Voxel-wise RSFA data from the (N=9,627) UK Biobank sample was averaged within each of 400 roughly symmetric volumetric ROIs from the 7-Network cortical parcellation of Schaefer and colleagues^47^. Parcel-wise RSFA values were residualized for the effect of age, sex, age^2^, ageXsex, age^2^Xsex, height, weight, BMI, combined gray/white matter volume (normed for head size), combined ventricular/CSF volume (normed for head size), diastolic and systolic blood pressure, run-wise rsfMRI motion, rsfMRI inverse SNR, T1 inverse SNR, and UK Biobank assessment center. Hierarchical clustering of residualized RSFA estimates was conducted using R in order to group regions with similar between-subject patterns of covariation. A 6-parcel RSFA clustering solution was selected. Raw RSFA values were then averaged across parcels falling within the same data-derived between-subjects cluster for use in heritability analyses. SNP-wise heritability of RSFA was estimated with genotyped data using GCTA-REML software. Age, sex, age^2^, height, weight, BMI, combined normed gray/white matter volume, combined normed ventricular/CSF volume, diastolic and systolic blood pressure, run-wise rsfMRI motion, rsfMRI inverse SNR, T1 inverse SNR, UK Biobank assessment center, and 10 genetic principal components were included as covariates.

Partitioned heritability analyses were conducted for the six RSFA clusters and for each of the 400 individual cortical parcels. Using AHBA expression data, genes were rank ordered by their spatial cortical correlation to *SST* and *PVALB*. Genes without Entrez IDs were removed. The BioMart package^90^ was used to identify each gene’s transcription start and stop sites (±5000 base pairs) according to the GRCh37-hg19 genome assembly. If no UKB genotyped variants fell within the intragenic regions of a particular gene, that gene was excluded from analyses. Otherwise, the gene was cross-referenced to cortical eQTL databases from the NIH GTEx project^55^ and CommonMind consortium^54^. Intragenic (±5000 base pairs) and eQTL SNPs associated with the top 500 *SST* (N_SNP_=8,612) and *PVALB* (N_SNP_=9,819) correlated genes were used for partitioned heritability analyses, respectively denoted *SST*SNP and *PVALB*SNP. Genetic-relatedness matrices for the *SST*SNP and *PVALB*SNP partitions were generated, as well as one for all remaining genotyped SNPs. RSFA heritability accounted for by each genetic relatedness matrix was estimated simultaneously for each of the three partitions using GCTA^49^. Partitioned heritability was then defined as the ratio phenotypic variance explained by either the *SST*SNP or *PVALB*SNP, divided by the total phenotypic variance. To calculate the significance of individual partitions, we consider the Wald test statistic against the null of 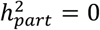, which follows a half-half mixture of 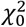 (a 𝒳^2^distribution with a probability mass at zero) and 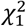 (a 𝒳^2^ distribution with 1 degree of freedom). Enrichment values were calculated to determine if the proportion of variability explained by a partition was greater than the proportion of variants within the partition, defined as

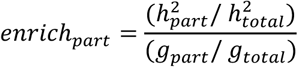

where 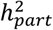 is the heritable variance explained by the SNP partition (e.g. *PVALB*SNP), 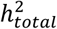 is the heritable variance explained by all partitions, *g*_part_ is the number of variants within the SNP partition, and *g*_total_ is the total number of genotyped SNPs. Standard error for SNP partitions were similarly scaled by the genome partition denominator. When calculating RSFA partitioned heritability across individual parcels (i.e. Figure 5), parcels with outlier partitioned heritability (i.e. *PVALB*_PART_, *SST*_PART_) and expression (i.e. *PVALB, SST*) greater than 4 standard deviations from the mean were excluding, resulting in 328 observations across cortex.

To assess whether schizophrenia polygenic risk was enriched among *SST* and *PVALB* correlated gene sets, competitive gene-set analysis was conducted using MAGMA^59^. Rank-ordered *SST* and *PVALB* genes were divided into twenty non-overlapping 500-gene bins. Schizophrenia summary statistics from the GWAS of Ripke and colleagues was used^58^. Intragenic variants were defined using a ±5000 base pair window, and gene set enrichment was estimated simultaneously across all 40 gene bins, revealing whether a particular bin is more associated with polygenic risk for schizophrenia than all other genes. Polygenic risk for schizophrenia^58^ was calculated using PRSice^60^. Only the top-SNP from the major histocompatibility complex was used for generation of individual risk scores. Benjamani-Hochberg False-discovery rate correction was conducted separately for each GWAS p-value threshold examined (e.g. correction for 6 tests at the GWAS p<1.0 threshold).

### NIH Blueprint processing

Publicly available microarray data from six adult macaque primates (3 Female) were downloaded from the Gene Expression Omnibus website (https://www.ncbi.nlm.nih.gov/geo; accession number GSE31613). Expression values were converted from log10 to log2. Data from two macaques (1 Female) were excluded due to sparse sampling across cortex. Samples from the following 10 cortical regions were included in our analyses: OFC, ACC, medial temporal lobe, temporal area, DLPFC, A1C, S1C, M1C, V1, and V2. The *collapseRows* function^82^ was used in R to select the probe with the highest mean expression and ComBat was used to remove residual donor effects. *SST* and *PVALB* expression were mean and variance-normalized to reveal relative expression differences across cortex.

### BrainSpan processing

Publicly available RNAseq reads per kilobase per million (RPKM) data from the Brainspan atlas were used to characterize patterns of interneuron-marker gene expression across development. Cortical tissue samples were analyzed from early fetal [8-12 post-conception weeks (pcw), N=10, samples=88], early/mid fetal (13-21 pcw, N=10, samples=88), late fetal (24-37 pcw; N=5, samples=27), early infancy (4 months; N=3, samples=22), late infancy (10 months; N=1, samples=8), early childhood (1-4 yrs; N=5, samples=41), mid/late childhood (8-11 yrs; N=2, samples=30), adolescence (13-15 yrs; N=2, samples=14), and adulthood (18-40 yrs; N=8, samples=85) developmental. RNAseq probes without entrez IDs were excluded and duplicated probes were removed by selecting the probe with the highest mean expression. Data was log2 transformed and the effect of donor was removed separately for each age group using ComBat. Gene expression was then mean- and variance-normalized across cortical tissue samples separately for each developmental stage. When multiple ages were present in a development stage, age was included as a covariate in a linear regression predicting normalized *SST* expression from normalized *PVALB* expression.

## Code Availability

Code used for these analyses will be made available upon publication at the following url: https://github.com/HolmesLab/Anderson2019_interneuron

## SUPPLEMENTAL FIGURES

**Figure S1.**
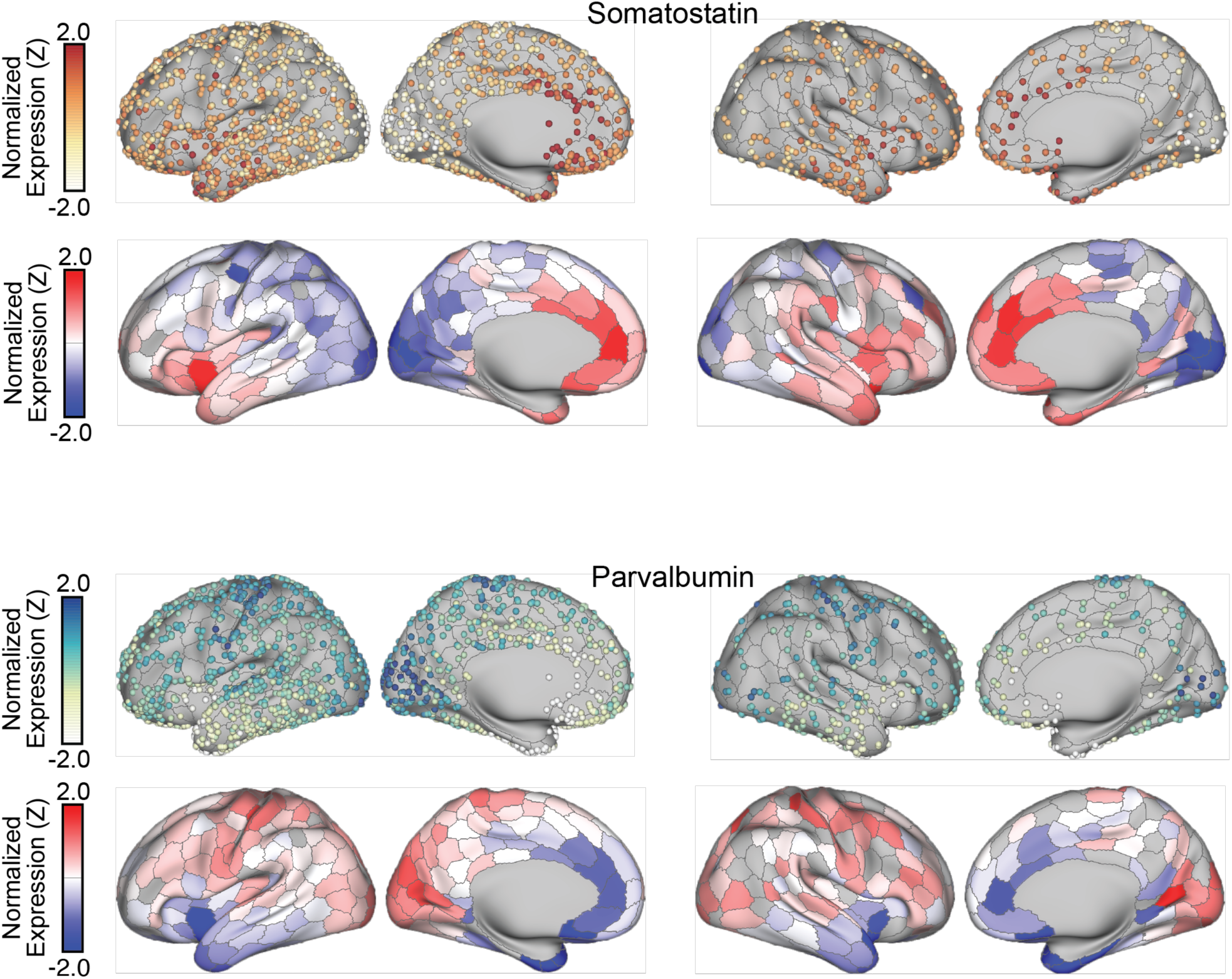
Sample and parcel-wise expression of interneuron markers *SST* and *PVALB*.

**Figure S2.**
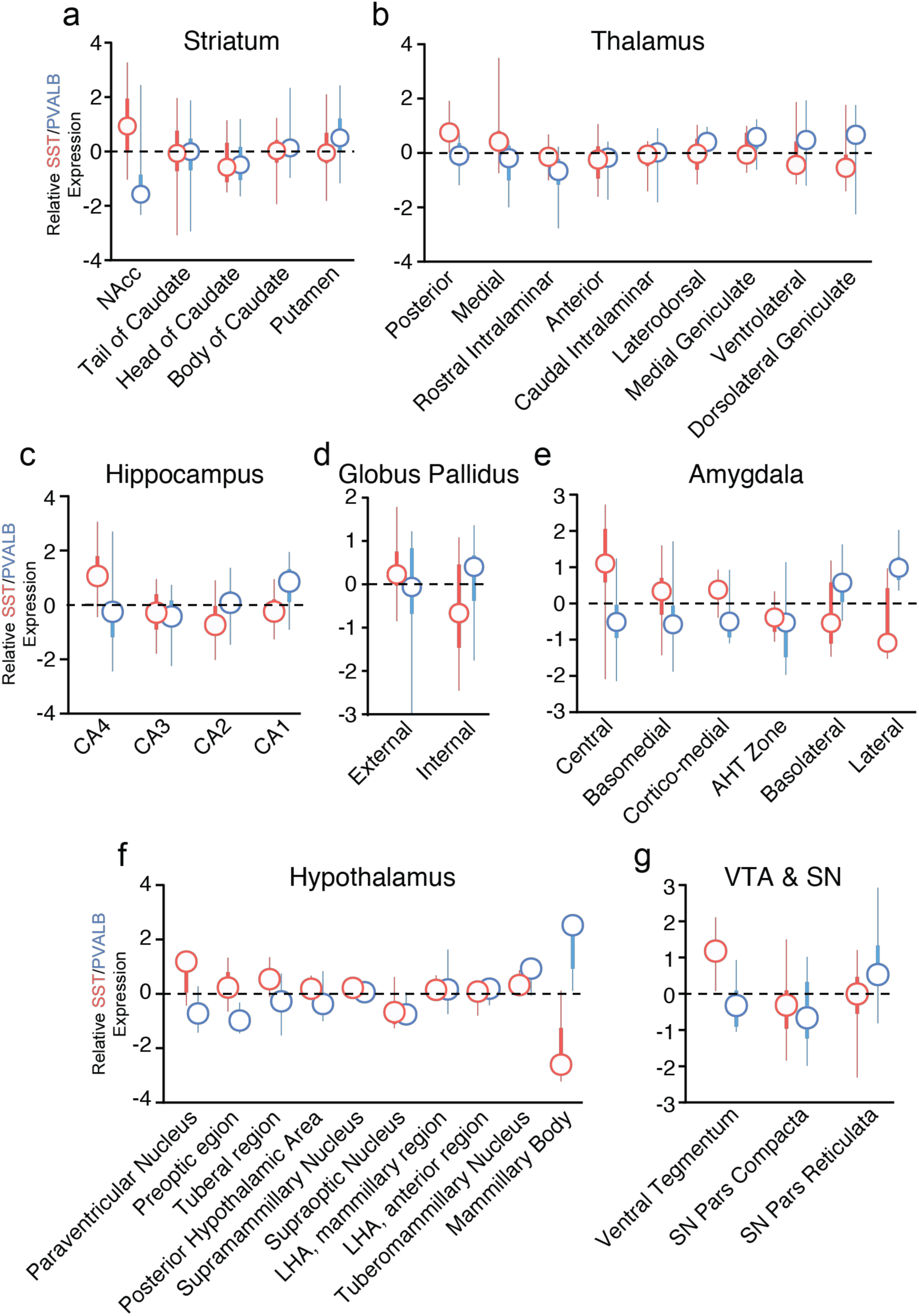
Z-transformed expression values of SST and PVALB for subregions of (a) striatum, (b) thalamus, (c) hippocampus, (d) globus pallidus, (e) amygdala, (f) hypothalamus, and (g) combined ventral tegmentum and substantia nigra. Regions are ordered by relative median expression of SST to PVALB. Circle=median, thick lines=interquartile range, thin line=minimum and maximum.

**Figure S3.**
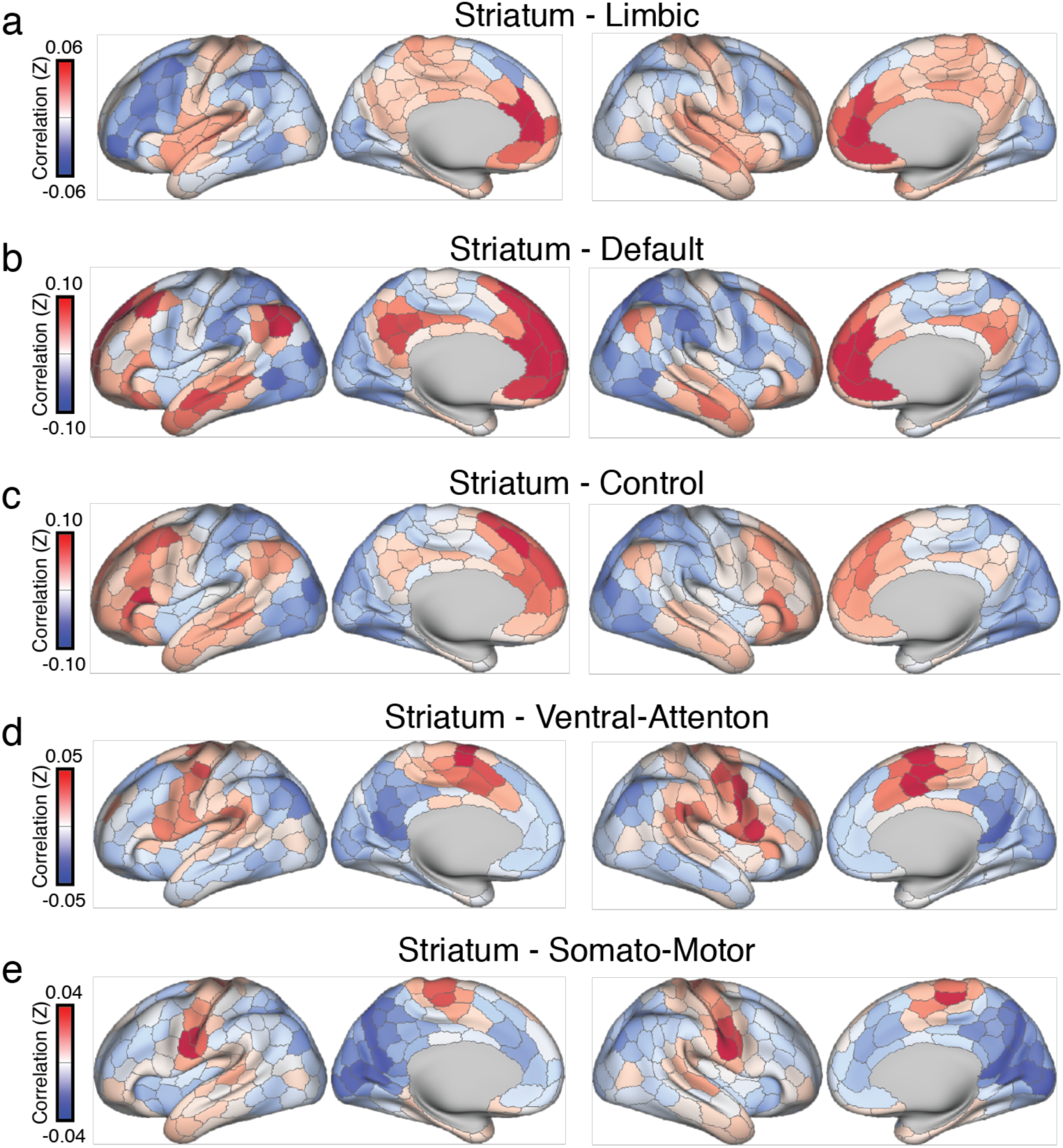
Cortical resting-state functional correlation profile for each of the 5 striatal parcels^39^ with analyzable AHBA expression data. Averaged maps were calculated using data from 9,627 participants from the UK Biobank.

**Figure S4.**
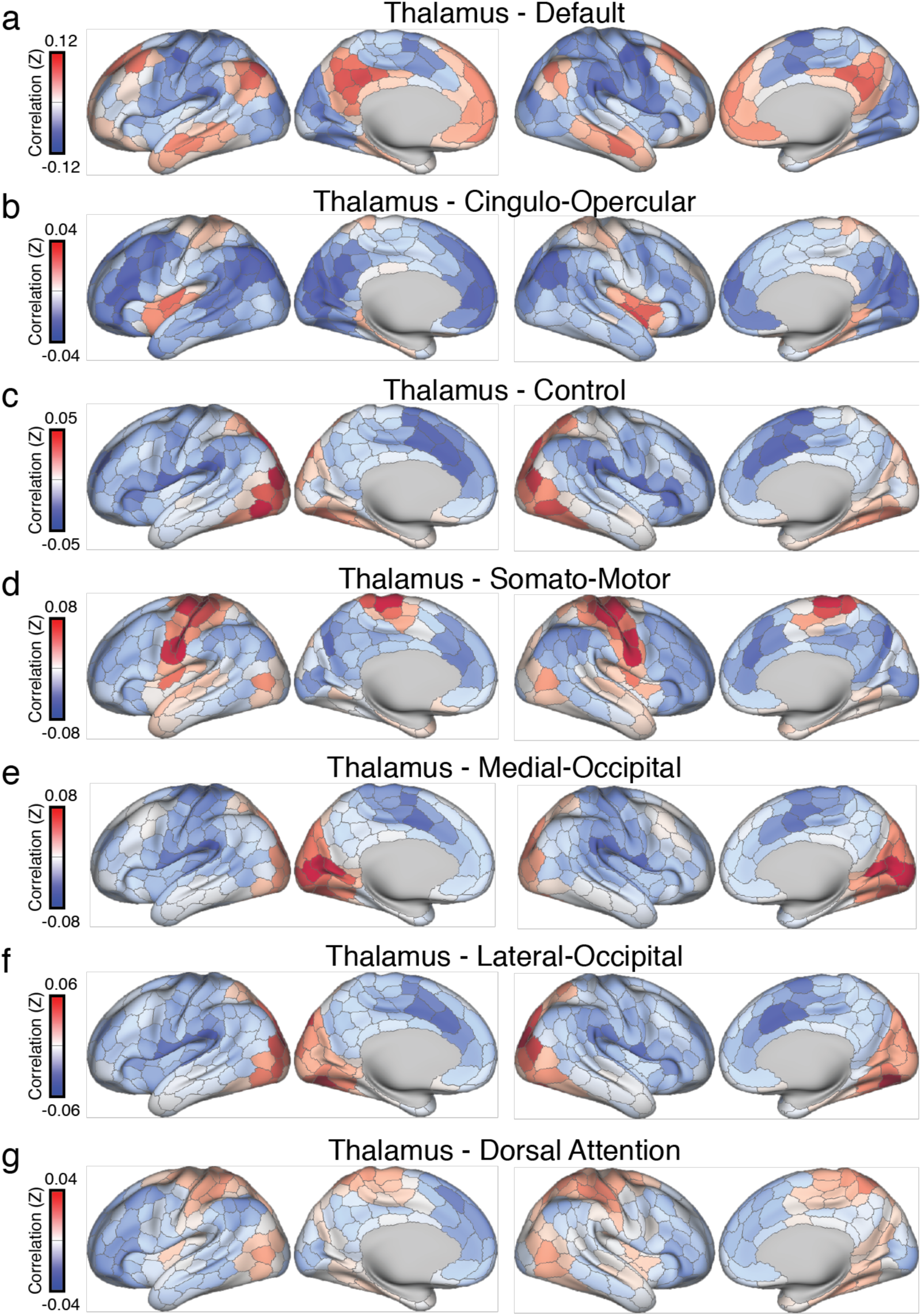
Cortical resting-state functional correlation profile for each of the 6 thalamic parcels with analyzable AHBA expression data^41^. Averaged maps were calculated using data from 9,627 participants from the UK Biobank.

**Figure S5.**
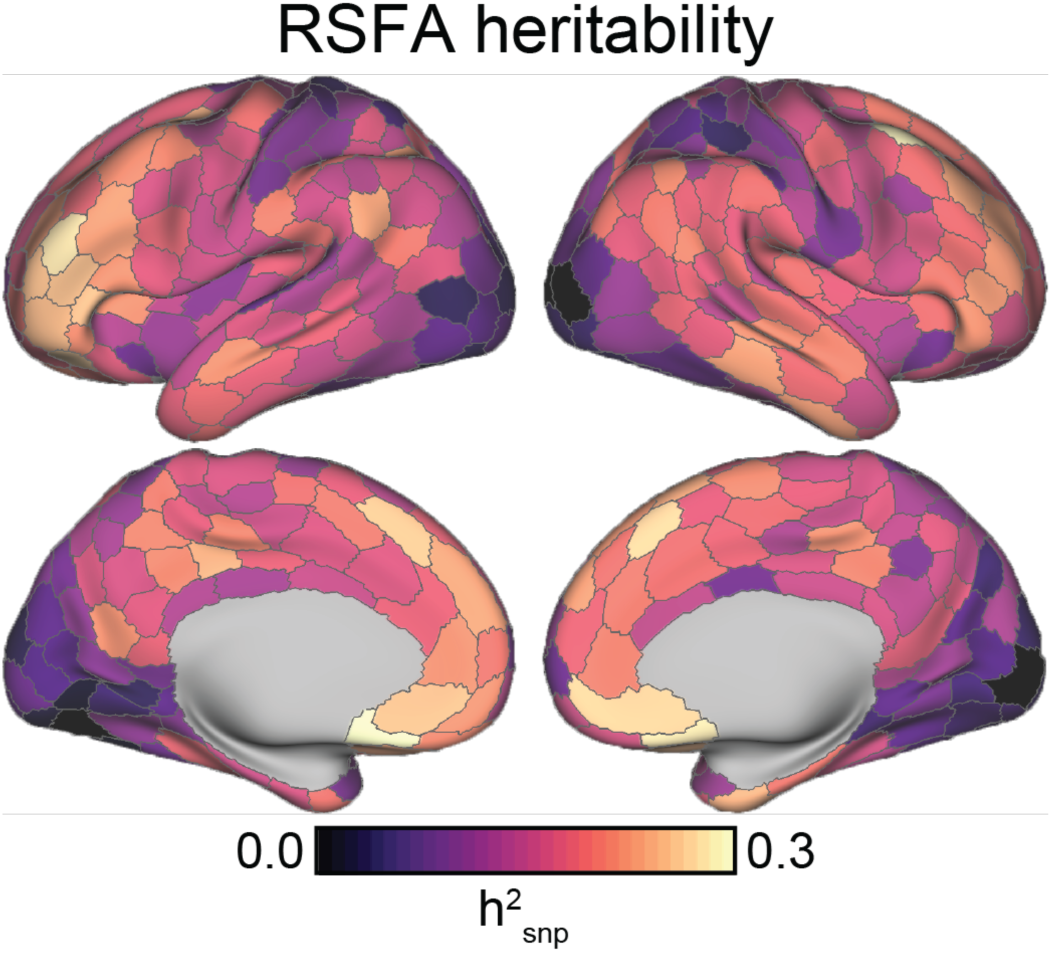
SNP-wise heritability for each of the 400 bi-hemispheric cortical parcels from Schaefer and colleagues^47^.

**Author contributions.** KMA and AJH designed the research. KMA conducted the research. MAC, RC, MDR, TG provided analytic support. KMA and AJH wrote the manuscript and made figures. All authors edited the manuscript.

## Acknowledgements

This work was supported by the National Institute of Mental Health (Grant K01MH099232 to A.J.H.), the National Science Foundation (DGE-1122492 to K.M.A.), and the National Institute on Aging (K99AG054573 to T.G.). Analyses were made possible by the high-performance computing facilities provided through the Yale Center for Research Computing. We thank B.J. Casey, Danielle Gerhard, Lauren Patrick, and Erica Ho for their feedback on early versions of the project. This work used data from the Allen Institute for Brain Science, NIH Blueprint Non-Human Primate Atlas, CommonMind Consortium, Brainspan Atlas of the Developing Human Brain, and has been conducted using the UK Biobank Resource under Application Number 25163. The Genotype-Tissue Expression (GTEx) Project was supported by the Common Fund of the Office of the Director of the National Institutes of Health, and by NCI, NHGRI, NHLBI, NIDA, NIMH, and NINDS. The data used for the analyses described in this manuscript were obtained from the GTEx Portal on 01/16/2018.

